# The feasibility and stability of large complex biological networks: a random matrix approach

**DOI:** 10.1101/223651

**Authors:** Lewi Stone

## Abstract

In his theoretical work of the 70’s, Robert May introduced a Random Matrix Theory (RMT) approach for studying the stability of large complex biological systems. Unlike the established paradigm, May demonstrated that complexity leads to instability in generic models of biological networks. The RMT approach has since similarly been applied in many disciplines. Central to the approach is the famous “circular law” that describes the eigenvalue distribution of an important class of random matrices. However the “circular law” generally does not apply for ecological and biological systems in which density-dependence (DD) operates. Here we directly determine the far more complicated eigenvalue distributions of complex DD systems. A simple mathematical solution falls out, that allows us to explore the connection between feasible systems (i.e., having all equilibrium populations positive) and stability. In particular, for these RMT systems, almost all feasible systems are stable. The degree of stability, or resilience, is shown to depend on the minimum equilibrium population, and not directly on factors such as network topology.

## Introduction

Network models have become indispensable tools for helping understand the biological processes responsible for the stability and sustainability of biological systems^1–18^. Intuitively, rich highly interconnected biological networks are expected to be the most stable, and are thus likely to better withstand the loss of a link, or to cope in the presence of external environmental perturbations. In the 70’s, May^1,2^ exploited random matrix theory (RMT), and the “circular law” for matrix eigenvalue distributions, to challenge this paradigm. He demonstrated that more complex and connected biological systems are in fact more fragile, and less likely to be stable, in terms of their ability to recover after some small external perturbation. Since then, the RMT framework has proved extremely useful for identifying those factors that beget stability in large ecological communities of randomly interacting species^5–15^. Moreover, in recent years, the modeling approach has successfully spread to other disciplines, ranging from systems biology, neurosciences, through to atomic physics, wireless, finance and banking, making this an exciting and vibrant contemporary research discipline^16–18^.

Here I re-examine similar issues of stability versus complexity, while using a better suited formulation of a biological system’s “community matrix” -- one that explicitly allows for the standard textbook assumption of density-dependent (DD) growth^2,19,20^. Such growth proves to be the rule rather than the exception for many biological processes, yet surprisingly, very little is known about their stability properties. In principal, May’s conclusions are not automatically translatable to DD systems. As we shall see, the “circular law” which sits at the foundation of May’s analysis, and governs the eigenvalue distribution of random matrices, generally does not hold for DD systems. The problem has resurfaced in recent prominent studies of ecological networks^6^.

In this paper we develop new methods to predict eigenvalue distributions of large complex DD systems. In the process, the analysis leads to and justifies new conclusions about the currently topical constraint of *feasibility*^6,7,10,15^. Feasibility requires that all equilibrium populations of a system are positive, a characteristic feature that is generally to be expected for any persistent system. There have been numerous reports in the literature of a strong association between feasibility of DD systems and stability, similar to Roberts (1974) who found that almost all feasible model systems are stable (see also Refs.10–12).

### Robert May’s model of large complex systems

It is helpful to first recall the original argument of May^1^. For an *n*-species community, let us suppose that the i’th species has abundance at time *t* given by 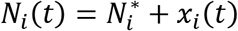. Here 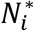 is the abundance at equilibrium (the symbol * indicating equilibrium), and *x_i_* (*t*) its perturbation from the equilibrium value. The dynamics of the populations are assumed to follow some complex nonlinear differential equation, which when linearized around equilibrium is of the form:

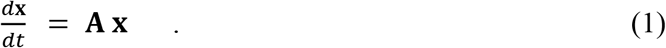

The vector **x** = (*x_i_*) contains the population disturbances *x_i_*, in terms of their perturbation from equilibrium, while the element *a_ij_* of the “community matrix” **A** represents the effect species-*j* has on the growth of species-*i* when close to equilibrium. A cooperative effect implies *a_ij_* > 0, while a negative effect is just the opposite with *a_ij_* < 0. The self-interactions between species are all scaled such that *a_ii_* = −1.

May^1^ studied communities under the limited *“neutral interaction”* assumption where interspecific interactions are equally positive as negative, and their expected or average value is zero i.e., E(*a_ij_*)=0. Environmental fluctuations are assumed to perturb the interaction strengths, so that for the basic “neutral interaction” model, **A = −I + B**, where **I** is the identity matrix and B is a random matrix with coefficients having mean zero and variance Var(*b_ij_*)=*σ*^2^. Finally, to model connectance of the interaction network, a proportion (1 − *C*) of randomly chosen interactions *a_ij_* are set to zero, leaving a proportion *C* nonzero.

More formally, we are interested in the “local stability” of biological models, which guarantees that a system will return to equilibrium after a “small” population perturbation. Unless otherwise stated, the paper will be concerned exclusively with local stability. A major achievement of May^1^ was to demonstrate that eqn.1 is locally stable for the neutral interaction model, if the interaction disturbances are “not too large,” namely if:

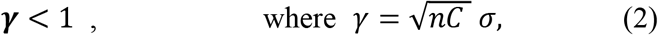

and unstable otherwise. The larger the number of species n, the sharper the transition from stability to instability at ***γ*** = 1. This is visualised in Fig.1a which plots the percentage of random matrices that are locally stable as a function of disturbance ***γ***. In terms of model parameters, the threshold criterion means that if either *n*, *σ* or *C* become too large, the system will transition into an unstable regime. With this simple but powerful argument, May demonstrated the fragility of large complex and highly connected systems.

**Fig 1.**
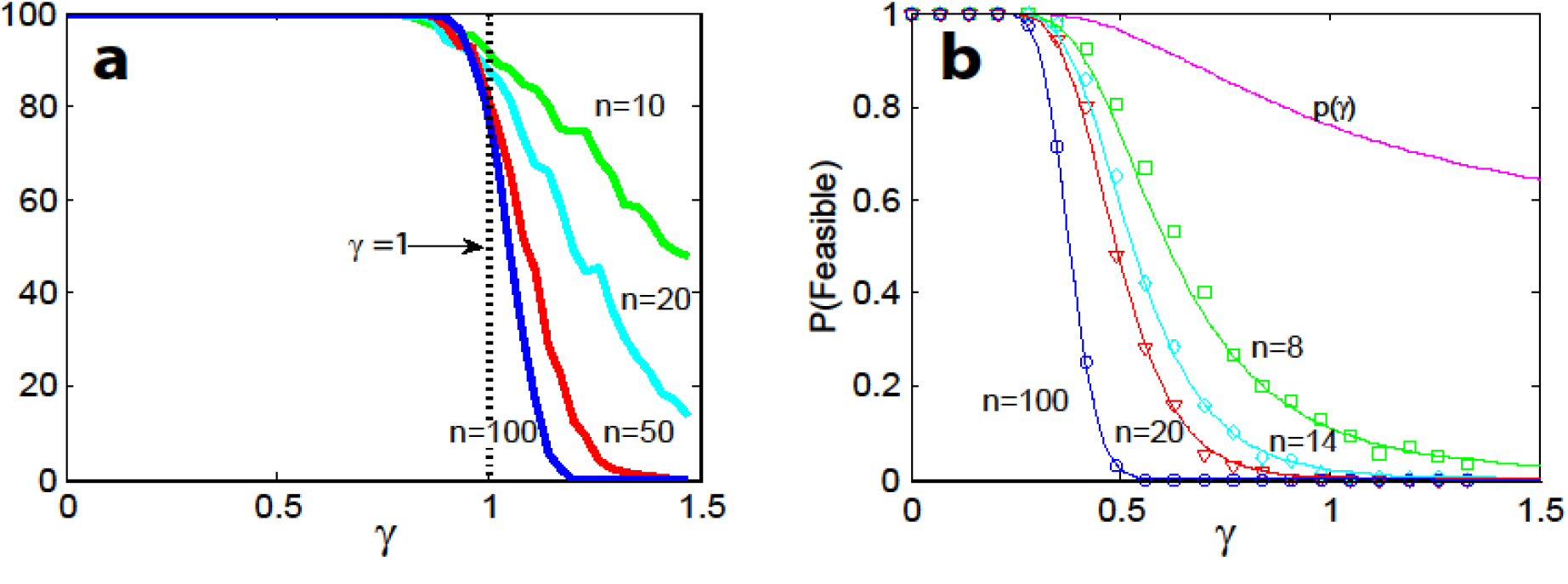
**a)** Percentage of locally stable interaction matrices **A** as a function of disturbance *γ* in an ensemble of 500 matrices for different community-sizes *n = 10,20,50,100*. May’s stability threshold sits at *γ* = 1. **b)**. The probability of feasibility, *P*(Feasible), as a function of disturbance *γ*, for n-species competition with different community sizes *n*=1,8,14,20,100. Each probability marked by a square, circle, etc is the proportion of feasible systems in 500 runs of eqn.9. Analytical predictions from eqn.12. Figure from Stone (1988, 2016).

## Results

### The eigenvalue distribution of the community matrix S = DA

Theoretical ecologists study the eigenvalues (*λ_i_*) of the community matrix **A** to determine local stability^2^. The theory underpinning the above elegant stability criterion relies on the “circular law” which is a central result in RMT. In simple terms, it states that the *n* eigenvalues *λ_i_* of the random matrix **A** are distributed uniformly in a circle with radius *γ* in the complex plane, and centred at (−1,0) on the real axis as shown in Fig.2a (see Ref.1). In this paper, it is often of interest to study the properties of each new matrix **A**, as ***γ*** is increased incrementally from zero. If the radius is increased to the point where it exceeds *γ* = 1, the eigenvalues of **A** populate the right-hand-side (RHS) of the complex plane indicating that at least one eigenvalue has a real part that is positive. The latter is the well known condition for triggering instability, and explains stability criterion eqn.2. In mathematical terminology, stability depends on the critical eigenvalue of the community matrix that has the largest real part, i.e.,

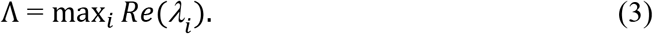

The system is locally stable iff Λ < 0, as in Fig.2a, since no eigenvalue has a positive real part.

**Fig 2.**
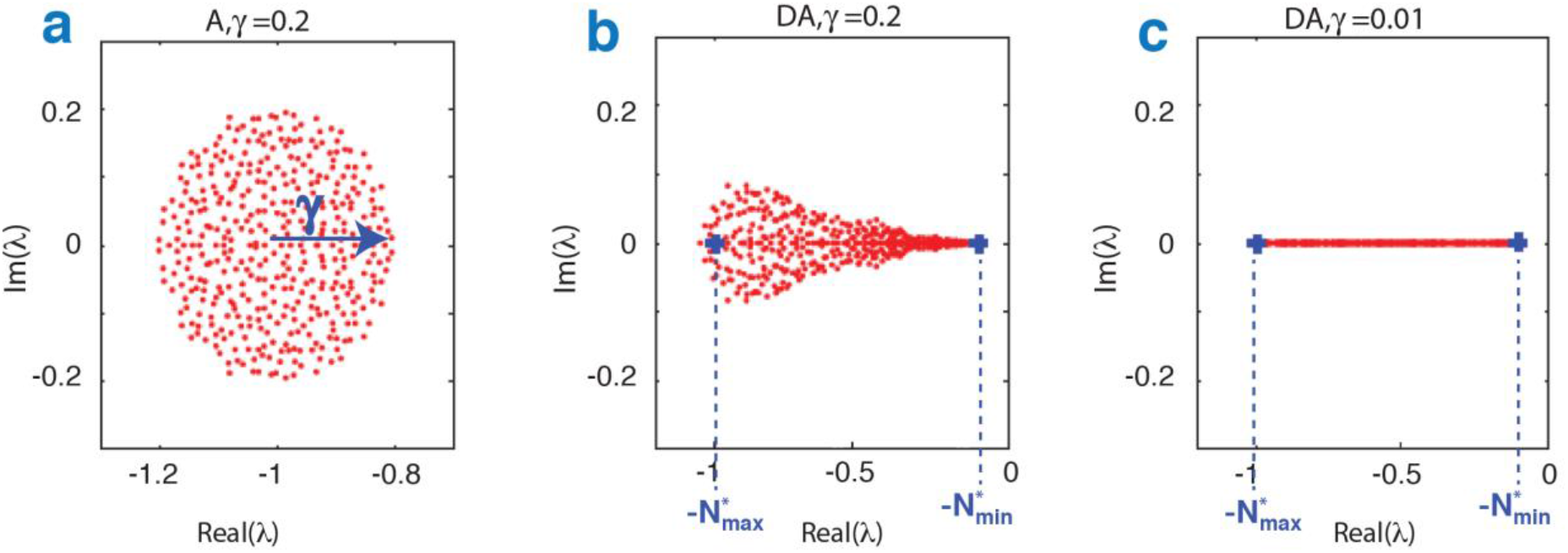
**a)** The distribution of eigenvalues of the matrix **A** in the complex plane for n=400, γ=0.2. Eigenvalues are distributed according to the “circular law” and fall in a circle centred at (-1,0) having radius γ (SI1). **b)** The eigenvalue distribution for the community matrix **S**=**DA**, where 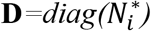 is a positive diagonal matrix with the same matrix **A** as in (**a)**. The circular distribution disappears and is replaced by a “guitar-shaped” distribution in which the imaginary components of the eigenvalues appear flattened out compared with (**a)**. The extreme left-hand and right-hand eigenvalues are predicted well by 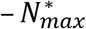 and 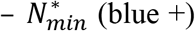. (c) Same as (b) but with γ=0.01. Now nearly all eigenvalues are real and sit close to the real axis wedged between 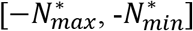.

More plausible biological models that include the operation of density-dependence (DD), may be framed in the form

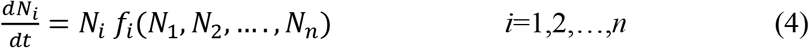

which provides a general characterization of many biological systems^19–21^. In these models, the net per-capita growth rate of an *individual* of species-i, depends on its interactions with other species as defined by some (often complicated) function 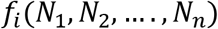. In the simplest DD model, 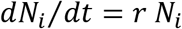, and each species has the same constant per-capita growth rate r, giving rise to exponential growth for each species. The well known Lotka-Volterra equations are a paradigmatic example of a more complex DD model, as discussed below.

We will be interested in determining whether or not a feasible equilibrium solution of eqn.(4), 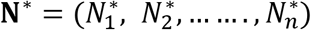, is stable. But stability of this equilibrium is no longer solely determined by matrices of the form **A**, as defined earlier. Instead, stability is determined by the critical eigenvalue of the community matrix **S = DA**, where the diagonal matrix 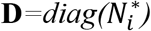^2,19,20^. The matrix **A** is the usual ecological species interaction matrix, whereby *a_ij_* represents the *per capita* effect species-*j* has on the growth of an individual of species-*i* (see Ref.21). Again, local stability of a feasible equilibrium is guaranteed iff the critical eigenvalue of **S** satisfies Λ < 0. It is important to emphasise, that even though the matrix **A** might be stable, this doesn’t automatically imply the matrix **S=DA** is stable (**D**>0), and this can potentially create problems^6–10^.

To see this in practice, a useful although hypothetical starting point is to assume that all *n*population equilibria 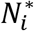 are randomly distributed in the interval (0,1), and then examine the community matrix **S=DA**, taking **A** as a random matrix. While **A** has eigenvalues distributed in a circle in the complex plane as shown in Fig.2a, this is no longer the case for the community matrix **S=DA** which now has a “guitar-shaped” distribution as seen in Fig.2b. The circular law for **A** becomes stretched and distorted as an outcome of the multiplication with the matrix of population densities 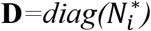.

Here we show how to extract the eigenvalue distribution for **S=DA**. In a recent important paper in the context of neuronal networks, Ahmadian et al. (2015; Ref. 22) studied the eigenvalue distribution of matrices having forms similar to the *n*x*n* stability matrix **S=DA**. If we set **A =−I + B**, where **B** is a mean zero random matrix, their results imply that for large *n*, the eigenvalue density of **S** is nonzero in the region of the complex plane, satisfying:

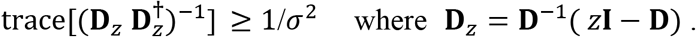

The complex variable *z* = *x* + *iy*, and trace 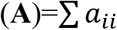 is defined as the usual sum of the matrix **A**’s diagonal elements. Applying the above inequality to the community matrix **S=DA**, it is not hard to show that the region corresponds to those values of *z* = *x* + *iy* for which:

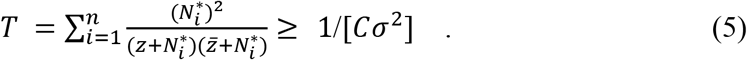

The inequality specifies a well-defined region in the complex plane where the eigenvalues of **S** lie. The region is referred to as the “support” of the eigenvalue distribution, and unlike the RMT circular law, the eigenvalue density is generally not uniform in this region.

The above inequality (5) shows that the support region of the eigenvalues is determined exclusively by the equilibrium populations 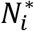, σ, and connectance *C*. Furthermore, one immediately notes that *T* has singularities at those points where 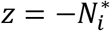, indicating that the region containing the eigenvalues of **S** must necessarily envelope the population equilibria 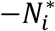. This gives an important hint of the strong relationship between the eigenvalues and the population equilibria.

It is possible to capture the complicated eigenvalue boundary that arises by evaluating eqn.5 at equality. Fig.3a plots the eigenvalue distribution for a typical community matrix S with *n=400, σ=0.01*, and *C=1* (i.e., *γ* = 0.2), while the 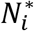 are chosen from a uniform distribution in the interval [0.05,1]. The boundary indicated in red is the curve deduced from eqn.5 evaluated at equality. Remarkably, eqn.5 accurately predicts the borders of the eigenvalue distribution, and that the red border envelopes all equilibrium populations: 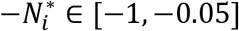.

Fig.3b plots the eigenvalues of **S** for both *γ* =0.2 (yellow dots) and *γ*= 0.9 (blue dots) superimposed on the same graph. Note that as *γ* is increased to *γ* = 0.9, the red boundary expands considerably. When *γ* > 1, the boundary moves into the RHS of the complex plane where eigenvalues have positive real parts (*Re*(*λ* > 0), and the system is unstable.

**Fig 3.**
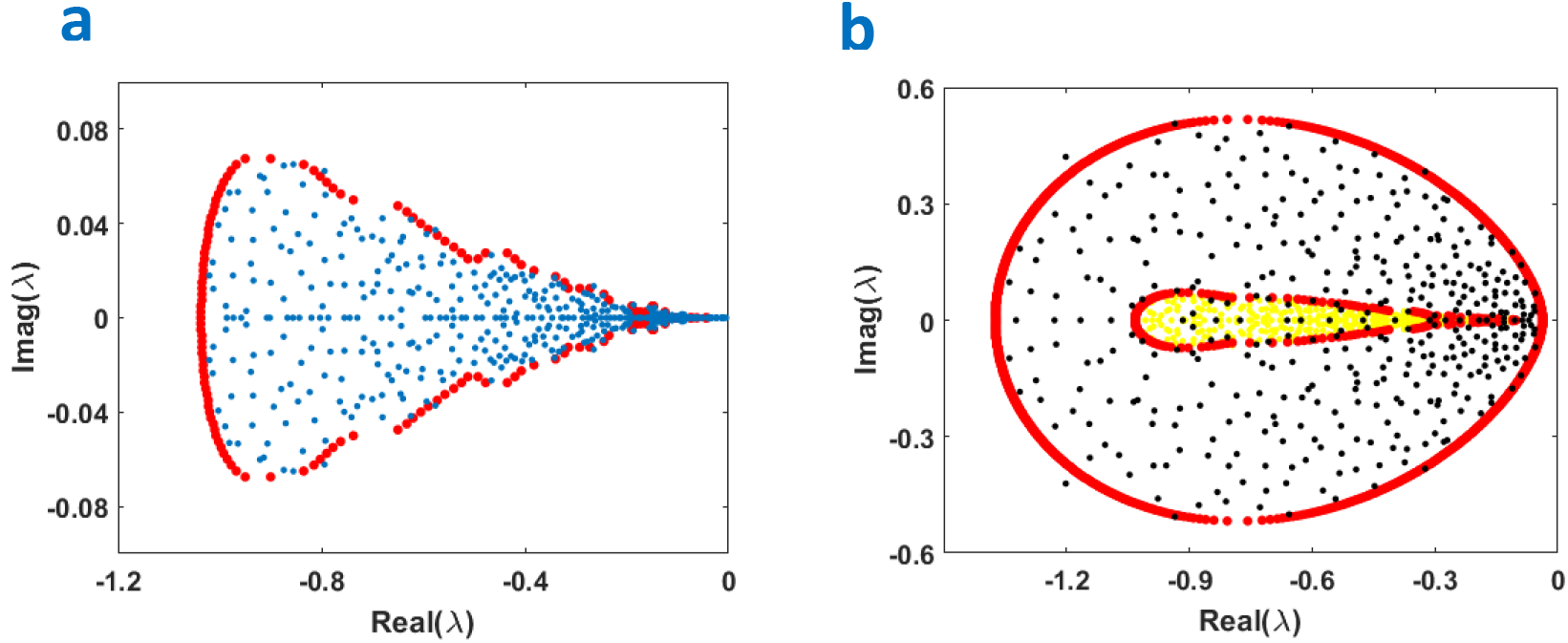
a) Eigenvalues (blue dots) of community matrix **S=DA** distributed in the complex plane, where= 400, *γ*= 0.2, 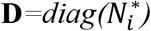, and 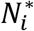 are uniformly drawn from interval [0.05,1]. Eigenvalue boundary appears as red dots, as obtained from eqn.5 evaluated at equality. **b)** Similar but with eigenvalues as yellow dots for *γ* = 0.2, and blue dots for *γ* = 0.9. The 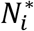 are uniformly drawn from interval [0.1,1].

Finally, it is not hard to see from eqn.5 that if we set all 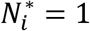, the May model is retrieved, and all eigenvalues lie in a circle in the complex plane centred at the point *z* = −1, having radius 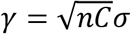. This of course retrieves the result indicated by eqn. 2, that stability is ensured if ***γ*** < 1.

For a given system, the eigenvalue distribution changes as 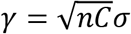 is increased, similar to that shown in Fig.3b for two different values of *γ*. For small *γ*, the eigenvalues all sit in the LHS of the complex plane, and the system is stable (the critical eigenvalue is Λ = max*_i_ Re*(*λ_i_*) > 0). As *γ* is increased beyond a threshold point, the eigenvalues start to populate the RHS and the system becomes unstable is ( A >0). For our situation, and based on eqn.5, as *γ* is increased from zero, the threshold between stability and instability occurs when the right-most point of the elliptical-like eigenvalue boundary (red line) first touches the point *z* = 0, or origin, in the complex plane. That is, where Λ = max*_i_ Re*(*λ_i_*) > 0). The threshold value of *γ*, can be found by evaluating eqn.5 at equality with *z*=0. This gives:

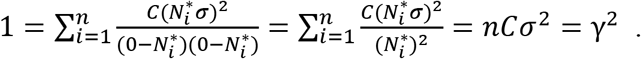

And thus the feasible equilibrium *N*\* is locally stable if:

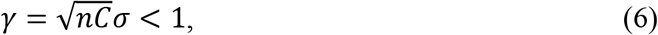

which surprisingly is independent of the positive equilibrium populations. The system is unstable if 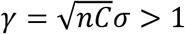. We thus find that May’s stability criterion is unusually general and holds for DD systems having community matrices of the form **S=DA**, even though the eigenvalue distributions of the latter are far from “circular.”

Note that the identical stability criteria (2) and (6) for A and **S=DA** are statistical criteria, and do not necessarily imply that the stability of the individual matrix A guarantees the stability of the matrix **S=DA**. However, based on the above results, it is demonstrated in SI2 that for these feasible RMT systems the matrices A and **S=DA** become unstable at exactly the same parameter values (approximately). That is, for a large feasible system (**D**>0),

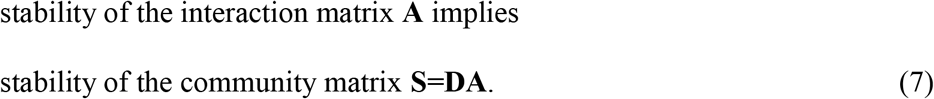

where **A** is a random matrix as defined by May^1^.

### Relationship between eigenvalues of S and the equilibrium abundances

Based on an “off-diagonal” matrix perturbation analysis it is possible to show that the eigenvalues of the community matrix **S=DA** of RMT systems and the equilibrium abundances 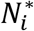, are simply related, namely: 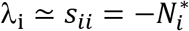 (see Methods and SI1). The approximation holds in the range *γ* < 1. Thus the critical eigenvalue component Λ = max*_i_ Re*(*λ_i_*), can be well approximated by the minimum equilibrium population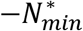:

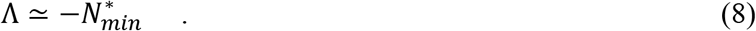

The critical eigenvalue component Λ = max*_i_ Re*(*λ_i_*), is often used as a stability or resiliency index^5,23,24^. When Λ is negative, the system is technically locally stable. However, the smaller or more negative is Λ, the more resilient is the community in terms of the time taken to return to equilibrium after a small perturbation. Arnoldi et al (2017) write that this form of “resilience is the most commonly used stability measure in theoretical ecology.” eqn.8 implies that the larger is the biomass of the rarest species 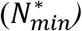, the stronger is the stability or resilience of the system since it will ensure a more negative, and faster return-time to equilibrium after perturbation^5,24^.

Since feasibility requires that the smallest equilibrium population 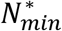 is positive 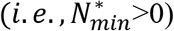, then eqn.8 makes transparent that in the regime *γ* < 1, feasibility is linked to both local stability (which requires Λ < 0) and resilience. Note that this result does not depend on any assumptions about randomness of the perturbation matrix **B**. To give an indication of the performance of eqn.8 as an estimator for, results for DD community matrices **S=DA** are given in Figs.4b and SI4. Some caveats and limitations concerning this approach are discussed in SI4&5.

## Implications and Biological Examples

### Example 1. Stability and eigenvalues of Lotka-Volterra competition communities

We now proceed to explore a fully defined density-dependent biological model, rather than just an abstract analysis of an arbitrary community matrix with random equilibria populations. The classical Lotka-Volterra (LV) equations serve this purpose well, being one of the most successful models for studying large complex systems^2–7,19^. For an *n*-species system, the equation for the abundance of species-*i* is:

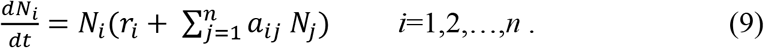

As before, the community matrix for this system may be written as the matrix **S=DA**, where now the populations in 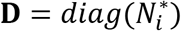 are actual equilibrium solutions of model eqn.9, found by setting all rates to zero. Following conventional practice, the intrinsic growth rates rare all scaled to unity^7,10,12,14^ (see SI1), with positive intrinsic growth rates r_i_ reflecting the implicit presence of resources. While some generality is lost with this scaling, it nevertheless opens the door to the advantageous possibility of analytical calculations.

The simplest competition community is the “uniform model,” where all coefficients are fixed to the same constant *a_ij_* = −*c*, 0 < *c* < 1, and the system is fully connected (*C*=1). In this parameter range, the equilibrium is always feasible and stable^11^. Hence the deterministic uniform model predicts that large competitive communities will satisfy two potentially advantageous features of viable ecosystems, namely feasibility and stability. We will see nevertheless that these seemingly stable and well-organized systems may be highly fragile in the presence of environmental fluctuations.

In the spirit of May (1972) and Roberts (1974), a large ensemble of competitive communities may be specified all of which, on the average, resemble the uniform model with mean interaction strength 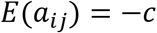. The interaction matrix **A** is given coefficients of the form 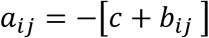, where the *b_ij_* are mean zero random perturbations with variance Var(*b_ij_*)=*σ*^2^.

Stability of the competition system depends in the usual way, on the eigenvalues of the community matrix **S=DA**. Fig.4a plots the eigenvalue distribution of the community matrix for a typical *n*=400 species competition community (*γ=0.3, c=0.1*) and we see that the red boundary for the support of the eigenvalues predicted by eqn.5 at equality, is an excellent fit. Note that the eigenvalues in Fig.4a are all in close vicinity, and are referred to as the “bulk” eigenvalues. The support region would appear to be even more contiguous if *n* were increased substantially. There is also an outlying real eigenvalue λ= -21.41 not shown in the figure as it is completely out of scale. For competition communities, the outlying eigenvalue which would sit at the extreme left of the complex plane, has no direct effect on stability. (It is in fact an outcome of having added a constant term-c to all interaction coefficients 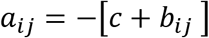.)

Because **S** has the properties of the “Google matrix,” it is shown in Refs.10&11 that for a competition system, all but one of the eigenvalues of the community matrix **S=DA** are *identical* (up to a scale factor of (1 − *c*)) to those for a system in which *c* = 0 (see Ref.25). But we have already worked out the stability properties when *c* = 0, via eqn.6&7. Hence, as shown in ^10–11^, it may be deduced by eqn.6 that:

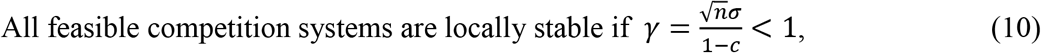

apart from rare statistical exceptions.

In Figure 4b, the real parts of the eigenvalues of **S** are plotted against the equilibrium populations 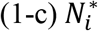 indicating 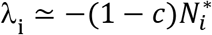, and the points sit close to the 45 degree line asjjredicted by the theory (eqn.8).

**Fig 4.**
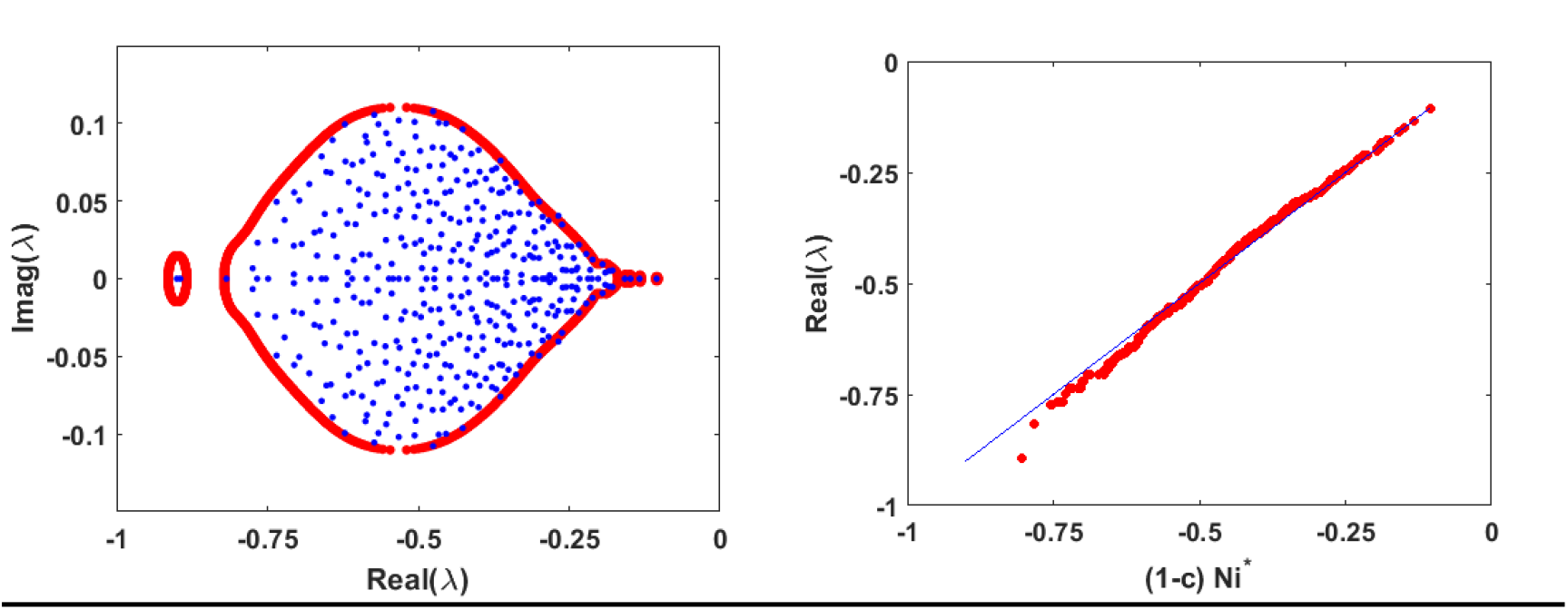
Competition community with *n=400*, *γ =0.3*, *c=0.1*. a) Boundary of the eigenvalue distribution (red) is plotted as predicted by eqn.5 and the actual numerically calculated eigenvalues are given (blue dots). b) The real parts of the eigenvalues of S are plotted against the equilibrium populations 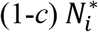 indicating 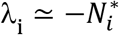, and the points sit close to the 45 degree line as predicted by eqn.8. For ease of visualisation the single outlying eigenvalue λ= -21.41 has been removed from the plots.

### Example 2. Feasibility implies stability in the ensemble LV model

We have just seen that when *γ* < 1 *all feasible systems* of the structure examined here are locally stable (apart from rare statistical exceptions). Without having gained an understanding of the eigenvalue relationship between the interaction matrix **A** and **S=DA** (see eqn.7), this result would not be available to us. An important question to ask now, is whether feasible RMT systems are always locally stable? This would be the case, if it could be shown that feasible systems only occur when *γ* γ 1.

To address this question, we determine the parameter regime where feasible systems can be found. The mathematical techniques required to accomplish this were presented in (Ref.10 and in Supplementary Information of Ref.11). The probability that a particular system is feasible *Pr*(Feasible) requires first the determination that a typical single species has positive population i.e., 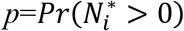.

#### Competition communities

Based on the equilibrium condition **AN^*^ =1** from eqn.9, when *γ* < 1 a first-order approximation of the equilibrium populations of the competition equations is

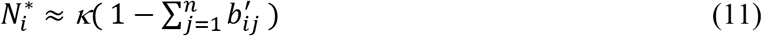

where *κ* is a positive constant and the symbol ′ represents a division by (1-*c*). This and higher order approximations, are discussed in Stone (1988, 2016-Supplementary Information).

[Comment: Not for publication. The following elaboration on calculating a probability of feasibility is based on Stone (1988 PhD thesis) and Stone (2016 in Supplementary Information). It has been added given critical readers of earlier drafts required clear direct evidence that feasible systems only exist for *γ* ≪ 1]

We let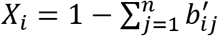 and note that by the Central Limit Theorem, *X_i_* is a normally distributed random variable with mean and variance:

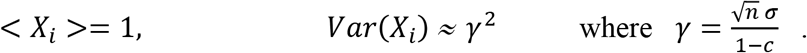

Thus 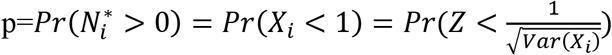, where Z is the standardized normal variate, namely *Z* ~ *N*(0,1)). Thus 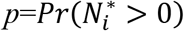 is purely a function of the single aggregated parameter *γ* i.e., *p* = *p*(*γ*).

Since the species are relatively independent, and since the *n*-species all have similar characteristics, a first order estimate of system feasibility is given by the probability that *all n*-species equilibria are greater than zero, namely:

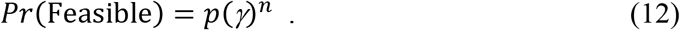

Fig.1b provides a plot of the percentage of feasible competition models from a random ensemble of 500 systems, as a function of disturbance *γ*. The graphs were generated for communities of different sizes from *n* = 14 to *n* = 100. Note carefully, that because of the exponent *n* in eqn.12, for large communities, feasible systems are only found for

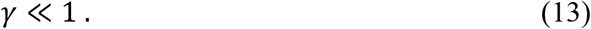

The graph shows clearly that the larger the number of species *n*, the more difficult it becomes to generate a feasible system. Analytical predictions based on eqn.12 are shown to be accurate in Fig.1b.

Return now to our initial query: Are all feasible systems stable? Results for competition communities indicate that feasibility is very fragile and strongly dependent on the variability in interaction strengths (i.e., *γ*) being sufficiently low; we should not expect to find feasible systems unless *γ* ≪ 1. Yet from Example 1 above, it was found that all feasible systems are stable as long as *γ* < 1. This implies *all feasible competition systems must be stable*. It also explains both in intuitive terms and theoretical terms (details in SI3) why there are no feasible-stable systems when *γ* < 1.

#### Mutualistic communities

For the case of mutualist systems, the result can be generalized further. Consider the LV *n*-species mutualistic system:

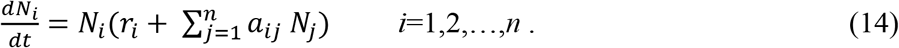

in which *a_ij_* ≥ 0, *a_ii_* = −1, and it is assumed the matrix **A** is strongly connected (i.e., irreducible). The birth rates *r_i_* ≥ 0 and at least one *r_i_* > 0. A simple application of M-matrix theory establishes that all feasible systems are locally stable^11,26^. More recently, this result has been extended and it has been shown that the mutualistic system Equations (13) possess a globally asymptotically stable feasible equilibrium iff **A** is locally stable^26^. This leads to an interesting situation with regards to mutualist systems, in that local stability of the interaction matrix **A** and feasibility are tied in a manner that ensures that *all feasible systems are stable*.

Constraints on feasibility for mutualistic systems are of a similar nature to those for competition (eqn.12), and demonstrate again that feasibility can only occur if *γ* ≫ 1.

### Example 3. Resiliency of competition versus mutualist communities

**A comparison of methods** The “great god of competition” concept^27^, has been a long-held principle amongst ecologists, for which competition is viewed as the main stabilizing force in ecological communities, while cooperation is viewed as essentially unstable. It is interesting to re-examine this principle by studying the stability properties of the interaction matrix **A**, and comparing with conclusions based on the community matrix **S=DA**.

#### i) Community matrix S=A, indicates mutualists are destabilizing

Based on the RMT system used by *(8)* suppose the Jacobian S is defined as

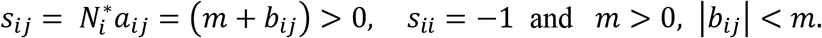

An underlying and unstated assumption is that all equilibrium populations are scaled to unity 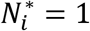, which effectively means we need only study the stability characteristics of the interaction matrix **A** with *s_ij_* = *a_ij_*. Thus from the outset, the framework assumes that a feasible equilibrium exists, which may be a wrong assumption.

The critical eigenvalue Λ = max*_i_ Re*(*λ_i_*) of **S=A**, for these mutualistic systems may be approximated (for large *n*) by the row sum of **S**, namely 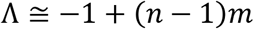. In short, Λ increases with *m*, and the feasible equilibrium becomes less and less resilient as *m* increases. A sufficient condition for instability of the equilibrium is that 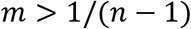, when the uniform model loses stability.

#### ii) Community matrix S=DA indicates mutualists have no effect or a positive effect on resilience

Using the more traditional approach of directly perturbing the species-interactions, as advocated here, the community matrix then has elements:

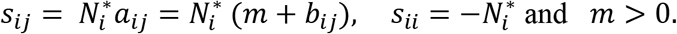

The equilibrium 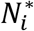 are solutions of the the LV model (eqns.9) whereby **AN*=−1**. Thus the community matrix **SN*= −1 N\***, has an eigenvalue of unity. This “outlier” eigenvalue is well separated from the “bulk” as shown in Fig.5. The critical eigenvalue of S proves to be Λ=— 1 for all values for which there is a feasible equilibrium. Thus the degree of mutualistic interaction *m* has *no* impact on the resiliency of a feasible equilibrium. If the average eigenvalue is used as an index to gauge resiliency of a feasible equilibrium, it is possible to show that the strength of mutualistic interactions *m* significantly increases the resiliency of feasible systems (see eg., Ref.26 and Fig.5).

Note however, that feasible stable mutualistic systems exists only for *m* γ 1 /(*n* − 1), i.e., only for relatively small communities having weak interactions. These communities can attain high population levels ensuring the community matrix **S=DA** has negative large magnitude eigenvalues (signals of strong stability). This very small parameter range for which feasibility is possible, does not indicate that mutualism is a highly unstable process. Outside of the narrow parameter range (i.e., for *m* < 1 /(*n* − 1)), no feasible equilibrium can even exist, so it makes no sense to discuss the stability or instability characteristics of an equilibrium that does not exist. The limited parameter range of coexistence, arises solely because of the difficult constraints in forming a feasible system, and has little relation to equilibrium stability. The mutualist model generates equilibria whose eigenvalues (real parts) become more and more negative, and thus stable, as the point of infeasibility is approached.

Competition systems, on the other hand, are stable if

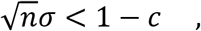

according to eqn.10 and based on the community matrix **S=DA**. Hence, competition appears destabilizing because increasing *c*, increases the critical eigenvalue 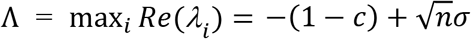, thereby decreasing resiliency.

The above analysis makes clear that the two different viewpoints **S=A** versus **S=DA** can lead to very different conclusions about resiliency. Analysis of the eigenvalues of the interaction matrix **S=A** supports classical competition theory, since it tends to show that ecological stability is enhanced by competition, while mutualism is highly destabilising. However, the study of the true community matrix **S=DA** (the method advocated here), finds mutualism should not be viewed as a destabilising process, and can often be stabilising in terms of resiliency, while competition is destabilising.

**Fig 5.**
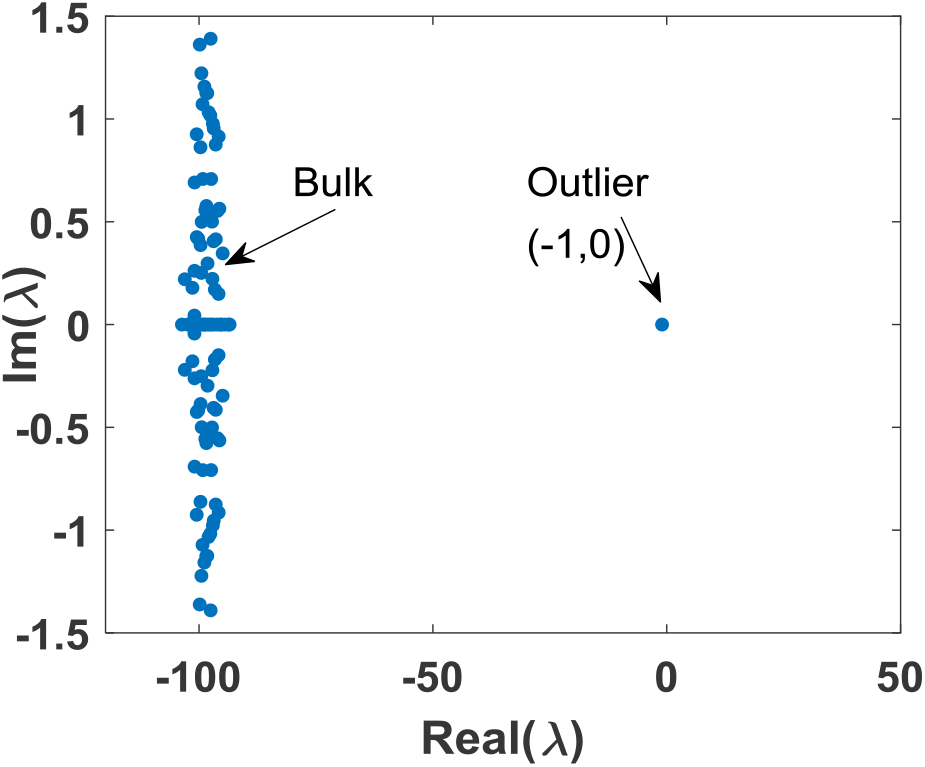
Eigenvalue distribution of community matrix **S=DA** for an n=100 species mutualistic community (m= 0.01, *σ*= 0.02. The stability of **S** depends on the critical outlier eigenvalue Λ = −1.

### Example 4. The impact of connectance *C* on feasible stable structured systems

Fig.6 examines the effect of connectance on predator-prey community matrices described in *(5,8)*. In these communities, each pair of species can only have signs indicative of predator-prey relationships of type (+,−) or (−,+), although the magnitudes of the interactions (*a_ij_*) are random and mean zero. Connectance *C* is included by incorporating a probability (1-*C*) that there is no interaction between the species whatsoever i.e., type (0,0). Two cases are examined:

i) the equilibrium populations 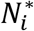 are given random values drawn uniformly in the interval (0,1) and kept fixed as connectance *C* is varied. Fig.6 plots both − Λ (green) and 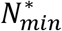 (black), and it is clear that 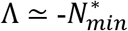, when *C* is varied over the full range, as predicted by eqn.8.
ii) The analysis is repeated for a Lotka-Volterra predator-prey model where the population equilibria are calculated from actual model parameters (*r_i_* = 1, in eqn.9). Again it is clear that the relationship 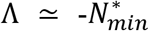 (red and blue respectively) holds, although now in contrast 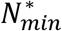 decreases as connectance *C* is increased.

Fig.6 clearly demonstrates that the two different models, random versus Lotka-Volterra, appear to respond to changes in connectance in qualitatively different ways, yet the underlying relationship 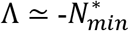, is preserved, and holds for both models. Thus the effects of connectance on system stability are highly model-dependent, and ultimately depends on how connectance affects the smallest equilibrium population. A related analysis of the impact of connectance on feasible competition systems is given in SI6.

**Fig 6.**
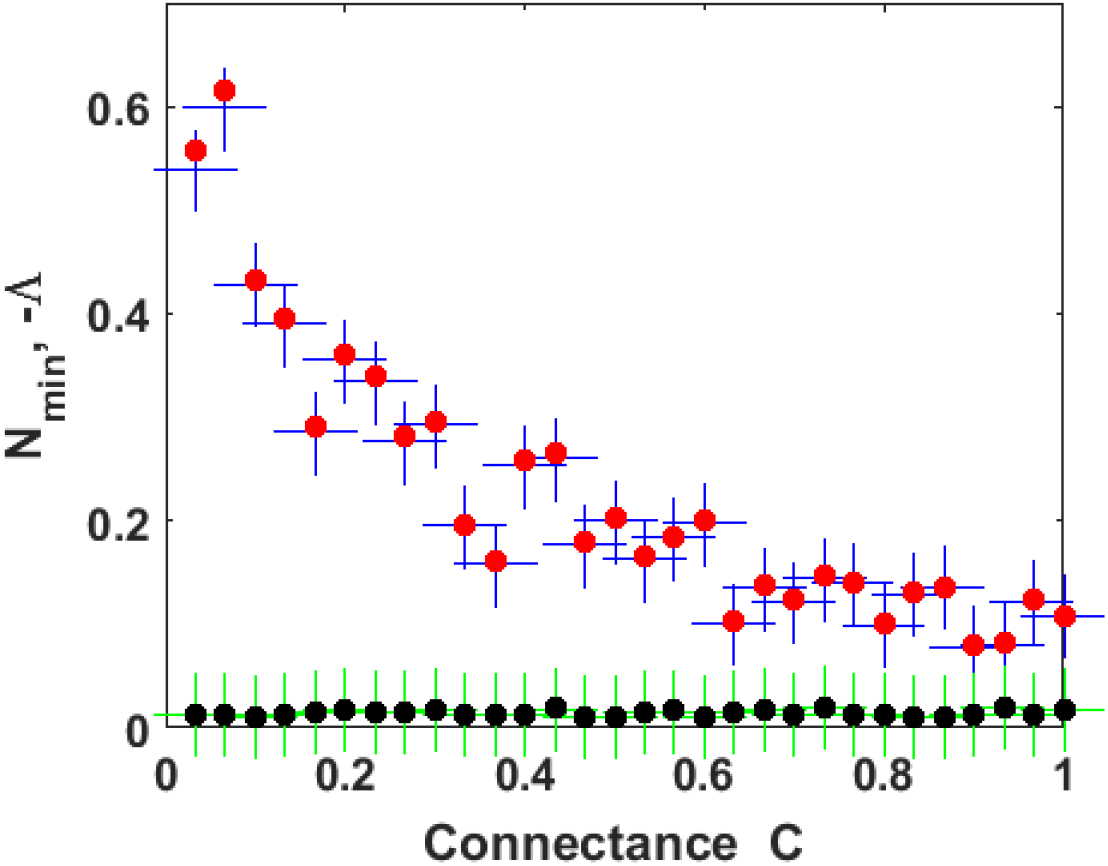
Predator-prey model *n*=200, γ=0.1. 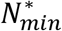 (red) and −Λ (blue) are plotted as a function of connectance *C*, as determined from LV equations (see text). A similar analysis, but with populations 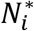 randomly chosen in the interval (0,1) 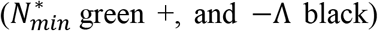. The graphs demonstrate 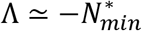 Nas predicted by eqn.8.

## Discussion

Many previous studies of biological networks have been unable to determine the stability properties of the community matrix **S=DA** for large complex random matrix systems. This is considered an unsolved and open problem^6,10^. Here a simple solution is presented based on the trace statistics of random matrices. For feasible RMT systems, it was shown that the community matrix **S=DA** transitions from stability to instability, at exactly the same parameter values for which the interaction matrix **A** transitions. Thus for a large feasible system with **D**>0, stability of the interaction matrix **A** implies stability of the community matrix **S=DA** (eqn.7).

Note that this is despite the fact that the matrix **A** has eigenvalues distributed according to the circular law while the eigenvalues of the community matrix **S=DA** are distributed completely differently (according to eqn.5). Because of the latter feature, the resilience characteristics obtained from analyses of the eigenvalues of **S=DA** are different to those obtained from the eigenvalues of **A**, as examples have demonstrated.

Feasible RMT systems were shown to be nearly always stable in the regime γ<1. However, for the classical ecological RMT models examined here, feasible systems are rarely found when γ>1. While these results may in some sense be model dependent, they should provide a good general characterization of how the addition of heterogeneity and external perturbations will affect any feasible stable system. Namely, as heterogeneity and disturbance increases, the feasibility of the system will be particularly sensitive to the heterogeneity in interaction, and feasibility will be lost often even before the transition from stability to instability of the interaction matrix. Future research in network science may benefit from shifting focus to study those factors which promote system feasibility^6,10,15^.

Finally, if the LV systems are a good guide to real world ecological systems, they inform us that large complex systems may be far more fragile than May’s main result predicts. The models studied here suggest that feasible stable mutualistic, competition and predator-prey systems can only be found if *γ* ≪ 1. Thus the models indicate the difficulty of assembling a large complex ecosystems that is feasible, and in addition indicate their fragility to perturbation in interaction strength. However, those RMT systems that can be assembled and are feasible, are nearly always found to be automatically stable. This helps explain why many large ecological networks *observed* in the real world (i.e., feasible systems) may be stable. It also suggests that large complex systems may be even more fragile than May’s main result predicts, and can be completely compromised when environmental perturbations exceed relatively small threshold levels. In the feasible regime, resilience of the most simple or the most complex network, is entirely dependent on the smallest equilibrium abundance, and not directly determined by network properties such as topology, modularity, clustering, and connectedness.

## Methods

### 1. Boundary of eigenvalue distributions

Ahmadian et al. (2015; Ref.22) studied the eigenvalue distribution of matrices having forms similar to the *n* × *n* matrix **S** = **M** + **DB**(where **M** and **D** are *n* × *n* deterministic matrices and **B** a random matrix), analysed here. They demonstrated^23^ that for large *n*, the eigenvalue density of the matrix **S** is nonzero in the region of the complex plane, satisfying:

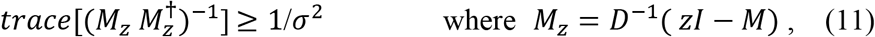

where the complex variable *z* = *x* + *iy*. This is equivalent to the region where:

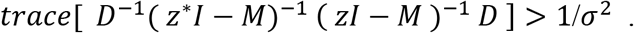

Thus the eigenvalue distribution is exclusively determined by the deterministic matrix **M**, in our case the equilibrium populations 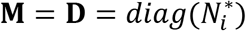, and the strength of random perturbations **σ**.

Assume now that **M= DQ**, where 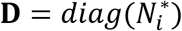 and **Q** is a symmetric matrix defining the topology of the deterministic component of species interactions. For May’s model **Q=I** is the identity matrix. In the case of the competition model 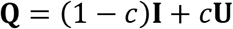, where all components of **U** are set to *U_ij_* = 1. To find the threshold between stability and instability for the matrix **S**, we note that the eigenvalues of **M** are real, and seek the point where they change from positive to negative as a parameter (eg σ or γ) is varied. We therefore seek the point where *z* = 0, and evaluate eqn.5 at equality to obtain:

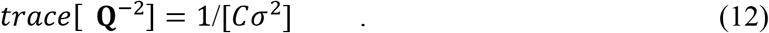

- For May’s system **Q=I**, and 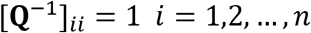 then, eqn.(12) becomes 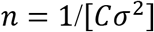, and we conclude the system is locally stable if 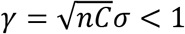.
- For the case of the fully connected (*C* = 1) competition system 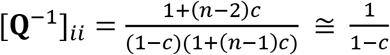 for large *n*. We thus conclude the system is locally stable if 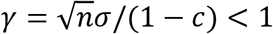. Here it is assumed that 0 ≤ *c* < 1.
- The criterion for more complex structured networks, can be determined by evaluating [*Q*^1^]_*ii*_ and then calculating the trace formula according to eqn.5.
- For the case of the fully connected mutualist systems,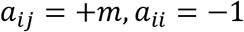, and 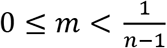, so that the “uniform” model is feasible. For feasible systems, the critical eigenvalue Λ = −1 is an outlier from the “bulk” of the eigenvalues.

### 2. Relation between population equilibria

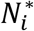 **and eigenvalues** *λ_i_*: Returning to inequality eqn.5, note that the left-hand-side of the expression for *T* has a singularity for those values of *z* for which 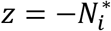. This is visualised in Fig.7 where *T* is plotted as a function of *z*, for an *n* = 10 species community with 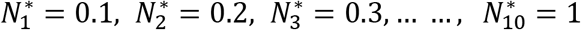. For purposes of illustration, it is assumed that *z* is a real number in the interval [0,1]. The function T clearly explodes at all points where 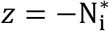. In this cut in the complex plane, the eigenvalues are predicted to be located on the x-axis (real-axis) at those points where *T* >1 / [*Cσ*^2^]=1000 (in this example). It is clear that the eigenvalues must lie close to the population equilibria 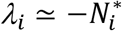. In general, the smaller the population 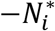 the more exacting is the approximation as can be seen from comparing the slopes of the graphs about the equilibria (and as can be verified by examining *∂T*/*∂x*).

This indicates that the smaller is σ, the closer the values of *z* are to the equilibrium solutions 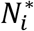, for which the inequality is satisfied. In fact, for σ≫ 1, we can expect 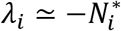. But this feature can already be observed in Figure 2, where it is clear that the eigenvalues in the complex plane are wedged between the vertical lines 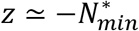 and 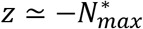.

### 3. The eigenvalue approximation

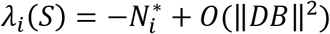

For the basic “neutral interaction” model, **A** = −**I** + **B**, where **I** is the identity matrix and **B** a matrix of perturbations that are not necessarily random. In the extreme limiting case, when all off-diagonal interspecific interactions are set to zero (*γ*=0), then **A** = −**I** and the community matrix is simply 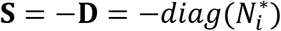, and the eigenvalues 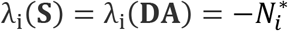.

When interspecific interactions are switched on (γ>0), and for reasonable assumptions (see also SI4 and [28]), the “off-diagonal” perturbation expansion is:

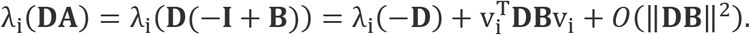

Here v_i_ is a normalised eigenvector of **D** such that 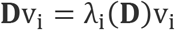. (The spectral norm 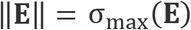 in terms of singular values may be used.) The success of the approximation eqn.8, is because the first-order perturbation term vanishes 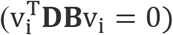, and

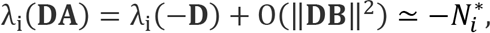

leaving a small quadratic error term (more details are given in SI4).

The intuition behind approximation eqn.8 may be understood as follows. In the extreme limiting case, when all off-diagonal interspecific interactions are set to zero (γ=0), the eigenvalues of **S=DA** have precisely the same magnitude as the equilibrium population values with 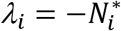, and therefore eqn.8 holds exactly. Fig.2c shows a situation very near to this case with γ=0.01, for which there are many weak off-diagonal interspecific interactions, and nearly all of the eigenvalues sit on the real axis in close proximity to the equilibrium population values 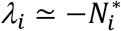. Denoting the smallest and largest equilibrium population as 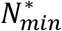 and 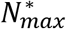 (in blue), then all eigenvalues should be wedged in the interval 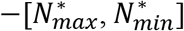 in the complex plane as seen in Fig.2c between the two demarked points in blue. But this holds to a good approximation even when the intensity of the perturbed interactions is increased, as shown for γ=0.2 in Fig.2b. See SI4 for more examples and a discussion of caveats regarding validity of approximation.

**Fig 7.**
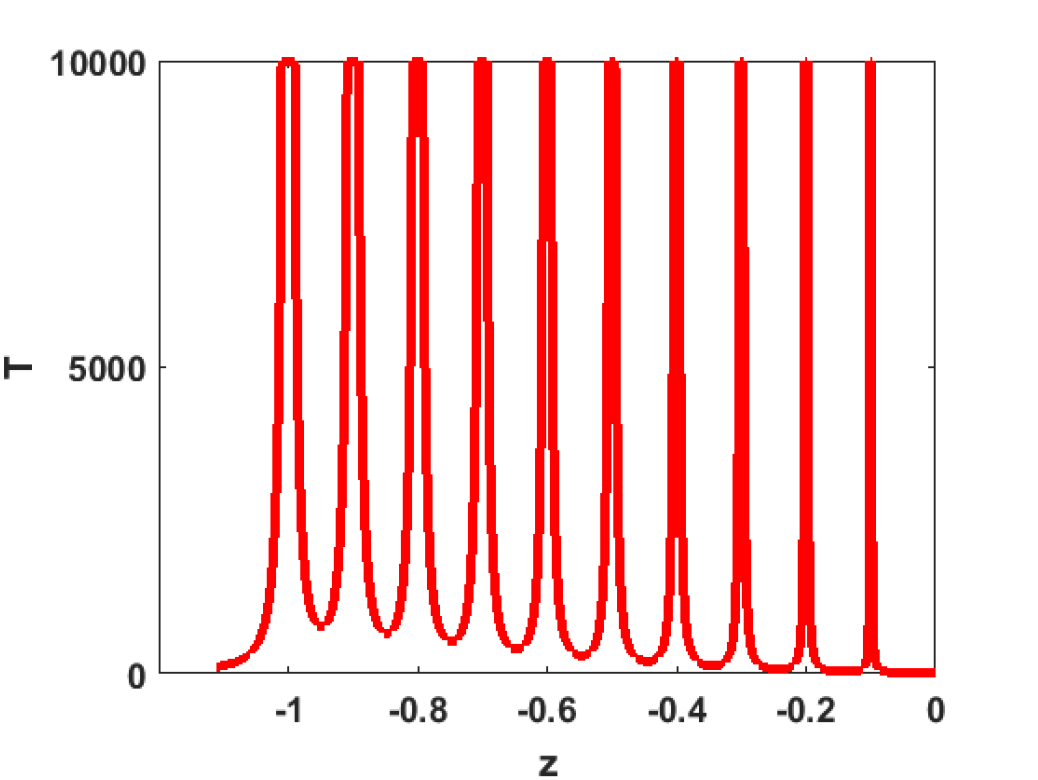
Plot of T in LHS of eqn.5 as a function of *x* = −*z*, which is real, for an *n* = 10 species community with 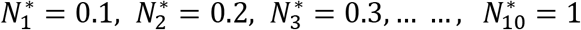. Singularities occur when 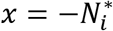.

## Acknowledgements

The support of the Australian Research Commission, for grants DP150102472 and DP170102303, is gratefully acknowledged.

## Author Contribution

All work carried out by Lewi Stone

## Data Availability Statement

No datasets were generated or analysed during the current study.

